# Role of the gut microbiota in the remodelling of the jejunal, colonic, hepatic and blood proteomes in a mouse model of Anorexia Nervosa

**DOI:** 10.64898/2026.10.28.762490

**Authors:** Jonathan Breton, Evy Timmerman, Lucie Houzard, Marion Huré, Colin Salaün, Delphi Van Haver, Jean-Luc do Rego, Marie Galmiche, Muriel Quillard, Marie-Pierre Tavolacci, Najate Achamrah, Pierre Déchelotte, Moïse Coëffier, Francis Impens, David Ribet

## Abstract

**Background:** Anorexia nervosa (AN) is a typical form of eating disorder characterized by a severe body weight loss, a fear of gaining weight and dysmorphophobia. This disorder is associated with both psychiatric comorbidities and multi-organ dysfunctions. Increasing evidence suggests a role of the gut microbiota in the onset or maintenance of AN. Although a dysbiosis has been reported in patients with AN, the physiological consequences of this dysbiosis remain poorly documented. To gain further insight into the role of the gut microbiota in the pathophysiology of AN, we used the murine Activity-based Anorexia (ABA) model and compared the remodelling of the jejunal, colonic, hepatic and blood proteomes occurring in male mice with intact gut microbiota, as well as in gut microbiota-depleted animals.

**Results:** We observed a significant proteome remodelling in several organs in response to the ABA model. More than 250 proteins involved in energy metabolism, catabolic processes, lipid metabolism, ghrelin regulation, and immune functions were found to show altered abundance in ABA mice compared to control mice. We observed in addition in the blood of ABA mice a significant increase in the level of circulating proteasome. By comparing mice treated or not with antibiotics, we identified 75 proteins whose regulation in ABA mice is gut-microbiota dependent. The other identified proteins were similarly affected in ABA mice with intact or depleted gut microbiota, suggesting that these factors are mainly regulated by food starvation and physical activity, and not directly by the gut microbiota. We finally confirmed that ORM2, a protein identified as significantly decreased in the plasma of ABA mice, is also decreased in the blood of patients suffering from AN.

**Conclusion:** Our results demonstrate that the Activity-Based Anorexia model is associated with an extensive proteome remodelling in several organs and tissues. We highlight in addition that some of these targets are regulated in a gut microbiota-dependent manner, thereby reinforcing the hypothesis of a role of the gut microbiota in Anorexia Nervosa. Together, our study provides a valuable resource to better delineate the impact of undernutrition and maladaptive physical activity on the proteome of various organs and the role of the gut bacteria in the pathophysiology of Anorexia Nervosa.

**Graphical abstract:**

- Male C57BL/6J mice were subjected to Activity-Based Anorexia (ABA) with or without gut microbiota depletion.
- Proteomic analyses were performed on jejunum, colon, liver, and plasma samples.
- Over 250 proteins are altered in ABA mice, which are mainly involved in catabolism, lipid metabolism, and immune pathways.
- A specific subset of proteins in the jejunum, colon, liver, and plasma is regulated in a microbiota-dependent manner in response to ABA, reinforcing the hypothesis that gut microbiota plays a role in Anorexia Nervosa.

## Introduction

Anorexia nervosa (AN) is a typical form of eating disorders (ED) characterized by a severe body weight loss, a fear of gaining weight and dysmorphophobia, according to the Diagnostic and Statistical Manual of Mental disorders (DSM-5) criteria. This eating disorder has an estimated prevalence of 1.4% for women and 0.2% for men and is considered as having the highest mortality rate among all mental illnesses^1,2^. AN represents a significant healthcare utilisation and an economic burden (annual healthcare costs per patient ranging from 3,000€ to 55,000€), which can be attributed to both psychiatric and somatic care^3^. In addition to restricting their food intake and/or engaging in binge-eating and purging behaviours that lead to a decrease in body weight, patients with AN exhibit multiple organ dysfunctions. These dysfunctions may affect the gastro-intestinal tract, with patients reporting abdominal pain, gastric hypersensitivity and constipation^4,5^. The may also affect the liver, with the occurrence of hepatic complications (elevated aminotransferases and hepatitis)^6^, as well as changes in blood parameters, such as an increase in the level of pro-inflammatory cytokines^7^. Altogether, these symptoms worsen patient’s condition and may further complicate refeeding strategies^4,5^.

Several studies have explored the potential role of the gut microbiota in the onset and/or maintenance of ED including AN^8–15^. The bacteria that make up the gut microbiota perform many functions and regulate a wide range of physiological processes, such as fat storage, energy homeostasis, or even host behaviour, via the production of diversified metabolites and proteins^16^. This symbiotic relationship with the host, leads to define the gut microbiota as a “metabolic organ”, which interact with various host organs via specific communication axes such as the microbiota-gut-brain axis or the microbiota-gut-liver axis. Several clinical studies have reported a gut dysbiosis in AN^12,13,17–28^. The features of this dysbiosis are not conserved between studies, which may be due to heterogeneity in clinical phenotypes and disease severity, or to experimental methodologies. Nevertheless, several bacterial taxa were repeatedly observed as increased or decreased in AN patients. *Methanobrevibacter smithii* for example, have been found overrepresented in AN patients^8,13,17,18,21,29^, while SCFA (Short Chain Fatty Acid)-producing taxa such as *Agathobacter, Romboutsia* or *Roseburia* were found underrepresented in several cohorts of AN patients^8,13,15,22,26,27^. Pathobionts or pathogenic bacteria were also observed as increased in AN patients compared to healthy individuals^15,19^. The exact role of the gut dysbiosis in AN physiopathology remains however unclear since identification of changes in gut microbial composition is not sufficient to establish causality.

Animal models of Anorexia Nervosa combined with advanced omics tools constitute valuable approaches to understand the pathophysiology of this eating disorder. The Activity-Based Anorexia (ABA) model is a useful rodent model to study AN, which combines both food restriction and physical activity via the use of activity wheels. This model is the most frequently used AN model, as it recapitulates numerous AN features including severe body weight loss, gastrointestinal disorders, in particular a delayed gastric emptying and an impairment of gut barrier functions, as well as a gut dysbiosis^30–36^

In this study, in order to better understand the potential role of the gut microbiota in AN, we compared the impact of the ABA model on the jejunal, colonic, hepatic and blood proteomes, in both control or gut microbiota-depleted animals.

## Material and Methods

### Animals

Animal experiments were approved by a regional Animal Experimentation Ethics Committee (authorization APAFIS#21102-2019061810387832). Experiments were carried out in accordance with current French and European regulations. Seven-weeks-old male C57Bl/6JRj mice (Janvier Labs, Le-Genest-Saint-Isle, France) were acclimated and housed at 23°C±2°C on reverse 12h light-dark cycle (dark phase from 10:00 am to 10:00 pm), with *ad libitum* access to water and food during 1 week. Animals were then randomly assigned to four experimental groups: a control group (CT group), a group treated with a cocktail of antibiotics in order to deplete their gut microbiota (CT-D group), a group with restricted access to food combined with a free access to a running wheel (Activity-based Anorexia group; ABA group) and an ABA group treated with antibiotics (ABA-D group).

### Antibiotics treatment

To deplete the gut microbiota of mice, CT-D and ABA-D mice had access to drinking water supplemented with a cocktail of antifungals and antibiotics, as previously described^37^. Briefly, 0.01 mg/mL Amphotericin-B (Sigma-Aldrich) was added in drinking water to prevent fungi overgrowth from day D_-7_ to D_-4_ of the protocol. From day D_-4_ to D_17_, drinking water was supplemented with 0.01 mg/mL Amphotericin-B, 0.5 mg/mL Vancomycin hydrochloride (Sigma-Aldrich), 1 mg/mL Neomycin trisulfate salt hydrate (Sigma-Aldrich), 1 mg/mL Metronidazole (Sigma-Aldrich) and 1 mg/mL Ampicillin (Sigma-Aldrich) (Fig. 1A).

**Figure 1:**
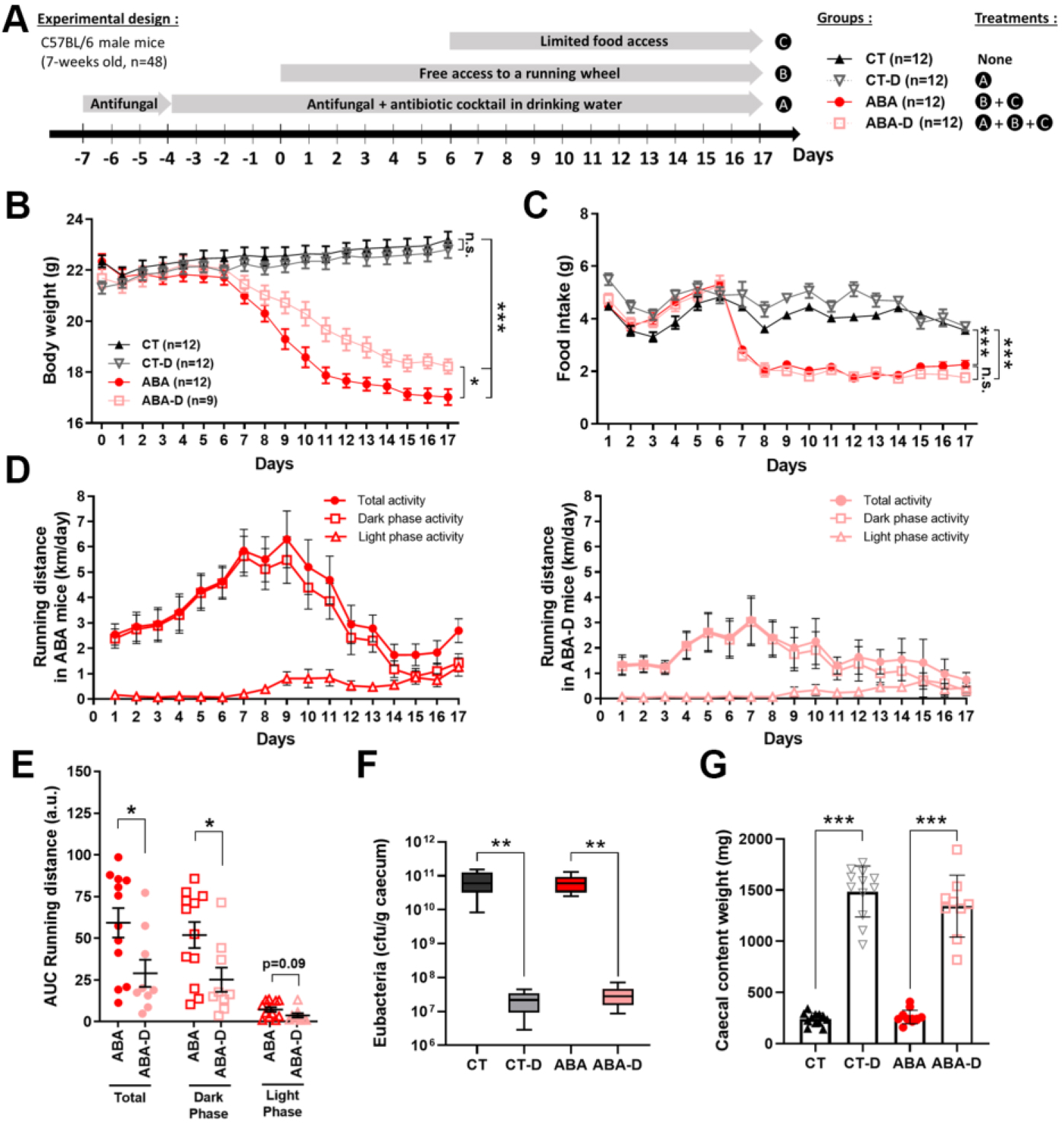
Changes in body weight and physical activity in response to the ABA model and to gut bacteria depletion. (A) Experimental design. (B) Body weight (means ± s.e.m.; n=9-12 per group; two-way ANOVA with Tukey’s correction; p-values at D_17_ are indicated. ABA vs CT and ABA-D vs CT, *** (p<0.001); ABA vs. ABA-D, * (p=0.048); CT vs. CT-D, not significant (n.s.; p=0.832)). (C) Food intake (means ± s.e.m.; n=9-12 per group; two-way ANOVA with Tukey’s correction; p-values at D_17_ are indicated; ABA vs CT and ABA-D vs CT, *** (p<0.001); ABA vs ABA-D, not significant (n.s.; p=0.212); CT vs CT-D, not significant (n.s.; p=0.915)). (D) Physical activity (means ± s.e.m.; n=12 for ABA mice and n=9 for ABA-D mice). (E) AUC (Area Under the Curve) of running distances (a.u., arbitrary units) (means ± s.e.m.; n=9-12 per group; Mann-Whitney; *, p<0,05; n.s., not significant). (F) Abundance of Eubacteria in caecal contents (values are represented as whisker plots; n=9-12 per group; one-way ANOVA with Tukey’s correction; **, p<0.01). (G) Caecal contents weight (means ± s.e.m.; n=9-12 per group; one-way ANOVA with Tukey’s correction; ***, p<0.001).

### Activity-based anorexia (ABA) model

The ABA procedure has been performed as previously described^32^. Briefly, the ABA model combines a free access to a running wheel combined with a progressively limited access to food from 6h/day at Day 6 (D_6_) to 3h/day at D_9_, and until the end of the protocol at D_17_. Food was given at the beginning of the dark phase (10:00 am). Animals have free access to drinking water (supplemented or not with antibiotics). Free access to activity wheels was provided to ABA and ABA-D mice from D_0_ to D_17_. The running wheel activity was continuously recorded during this period with the ActivityWheel® software (Intellibio, Seichamps, France). Food intake and body weight were measured daily at the end of the light phase. If body weight loss was higher than 20% on 3 consecutively days, animals were euthanized in accordance with the ethical procedures.

At the end of the procedure (D_17_), all animals were euthanized. Blood samples were collected with a standard 21 gauge needle, after puncturing the abdominal aorta, and transferred into BD Vacutainer K_2_EDTA tubes. These samples were then centrifugated twice (1,900 x g; 10 min; 4°C) to collect plasma and ensure to remove most platelets. 1 cm-long segments from the jejunum and colon were removed, immediately washed with ice-cold 1X PBS and frozen in liquid nitrogen. Caecal contents were collected in parallel, weighted and frozen in liquid nitrogen. Segments from the left lobe of animal’s liver were removed and frozen in liquid nitrogen. All samples were stored at −80°C until analysis.

### Evaluation of gut microbiota depletion induced by antibiotics

Total bacterial abundances were determined by quantitative real-time polymerase chain reaction (qPCR) on DNA extracted from mice feces collected at D_17_. Briefly, DNA were extracted using the QIAamp DNA Stool Mini Kit (QIAGEN), including a bead-beating step (0.1Vmm zirconia silica beads, BioSpec products, Bartlesville, USA)^34^. qPCR were performed using Itaq Universal SYBR Green Supermix (BioRad) and 16S rRNA primers targeting conserved regions in all Eubacteria (Eub-338 F: 5’-ACTCCTACGGGAGGCAGCAG-3’ and Eub-518 R:5’-ATTACCGCGGCTGCTGG-3’; annealing temp: 60°C)^37^. The Cq determined in each sample were compared with a standard curve made by diluting genomic DNA extracted from a pure culture of *E. coli,* for which cell counts were determined, prior to DNA isolation, by counting colony-forming units after plating.

### Proteomics analysis

A detailed description of the proteomics sample preparation, LC-MS/MS and data analysis is provided in the Supplementary Methods. Briefly, tissue and plasma samples were dissolved in a buffer containing 5% sodium dodecyl sulfate (SDS) and extracted proteins were digested with trypsin using the S-Trap protocol (Protifi). For tissue samples, the resulting peptides were TMT-labeled (TMTpro 16-plex, Thermo), pooled and pre-fractioned by RP-HPLC (Agilent series 1200). Pre-fractionated peptides were analyzed by LC-MS/MS analysis using 2-hour runs on an Ultimate 3000 RSLCnano system (Thermo) connected to a Q Exactive HF Biopharma mass spectrometer (Thermo) operating in data dependent acquisition (DDA) mode. Plasma samples were analyzed on the same instrument, but using an Evosep One (Evosep) with a 30 SPD method for peptide separation and data independent acquisition (DIA). Recorded spectra were searched against the mouse protein sequences in the Swiss-Prot database using MaxQuant for tissue and DIA-NN for plasma samples. STRING (version 12.0) and DAVID (DAVID 2021, DAVID Knowledgebase: v2023q4) were used as bioinformatic resources for functional annotation and enrichment analyses of the identified proteins.

### Clinical cohort and blood samplings

Patients over 18 years of age with clinician-diagnosed anorexia nervosa (AN), according to DSM-5 criteria, were recruited at the Nutrition Department of Rouen University Hospital, a care center specialized in eating disorders. These patients belong to the previously described EDILS cohort (Eating Disorders Inventory and Longitudinal Survey), and were selected based on the availability of plasma samples, clinical data, and gut microbiota composition data (Table 1)^15,38^. Healthy individuals with no active or past history of eating disorders and a Body Mass Index (BMI) between 18.5 and 24.9 kg/m² were recruited through the Clinical Investigation Center of Rouen University Hospital. All participants provided written informed consent. The study was conducted in accordance with institutional ethical guidelines was approved by the ethical committee N°CPP/CE 002/2014. Peripheral blood samples were collected in heparinized tubes from fasting participants during clinical evaluation. Samples were centrifuged and plasma aliquots were stored at −80 °C until analysis. Several questionnaires, including the Eating Disorder Inventory-2 (EDI-2), Hospital Anxiety and Depression Scale (HADS), Body Shape Questionnaire (BSQ), QUAVIAM, and the SF-36 questionnaire were filled by participants to get more insights into their personal traits and to detect potential comorbidities^15^. The relative abundance of the different fecal bacterial taxa was determined by performing 16S rRNA gene amplicon sequencing as previously described^15^.

**Table 1.** Clinical characteristics of the subset of participants from the EDILS cohort included in the present study.

| **Characteristic** | **AN (n = 34)** | **HV (n = 35)** | ***p*** **value** |
|---|---|---|---|
| **Age (years)** | 29.6 ± 11.3 | 38.3 ± 10.2 | **0.0002** |
| **BMI (kg/m<sup>2</sup>)** | 16.2 ± 1.9 | 21.8 ± 1.7 | **<0.0001** |
| **Female, n (%)** | 32 (94.1%) | 31 (88.6%) | 0.673 |
| **Male, n (%)** | 2 (5.9%) | 4 (11.4%) | — |
| **HADS depression score** | 7.7 ± 3.8 *(n = 32)* | 1.9 ± 2.1 | **<0.0001** |
| **HADS anxiety score** | 12.7 ± 3.8 *(n = 33)* | 5.5 ± 2.7 | **<0.0001** |
| **EDI-2 score** | 87.8 ± 37.8 *(n = 24)* | — | — |
| **BSQ score** | 95.9 ± 38.5 *(n = 29)* | — | — |
| **QUAVIAM score** | 292.5 ± 107.6 *(n = 28)* | — | — |
| **SF-36 PCS** | 70.5 ± 19.3 *(n = 22)* | 88.4 ± 11.4 | **0.0002** |
| **SF-36 MCS** | 55.8 ± 16.6 *(n = 21)* | 82.7 ± 11.5 | **<0.0001** |

Data are presented as mean ± SD. Sample sizes vary across variables because of missing data. EDI-2, BSQ and QUAVIAM scores were collected in patients with anorexia nervosa only. Comparisons between patients with anorexia nervosa (AN) and healthy volunteers (HV) were performed using the Mann–Whitney U test for continuous variables and Fisher's exact test for categorical variables. Statistically significant differences were considered at p < 0.05. AN, anorexia nervosa; HV, healthy volunteers; BMI, body mass index; HADS, Hospital Anxiety and Depression Scale; EDI-2, Eating Disorder Inventory-2; BSQ, Body Shape Questionnaire; QUAVIAM, QUAlité de VIe dans l'Anorexie Mentale; PCS, Physical Component Summary; MCS, Mental Component Summary; SF-36, SF-36-Item Short Form Health Survey.

### Protein quantification

Plasma levels of two candidate proteins, ORM2 (Orosomucoid-2) and PON1 (Serum paraoxonase/arylesterase 1), were quantified in plasma samples collected from participants, using commercially available ELISA kits according to the manufacturers’ instructions (ORM2; Elabscience, Cat. No. E-EL-H6086; PON1; Invitrogen, Cat. No. EH376RB). Briefly, standards and diluted plasma samples (1:50) were pipetted into the pre-coated microplate wells. After incubation and washing, a detection monoclonal antibody specific to human ORM2 or PON1 was added, followed by an HRP-conjugated secondary antibody. After the enzymatic reaction was stopped, color intensity was measured at 450 nm for PON1 and 425 nm for ORM2. ORM2 and PON1 concentrations in the unknown plasma samples were then determined from their respective calibrating curves.

### Statistical analysis

Animal body weight, food intake and physical activity were analysed using GraphPad Prism 8.3.0 software (GraphPad Software Inc., San Diego, CA, USA). Data were expressed as mean ± standard error to mean. Shapiro test was used to assess Gaussian distribution parameters. Data were analysed by one-way or two-way ANOVA, followed by Tukey posthoc tests. Comparisons of protein abundances between groups were performed using Perseus software (version 1.6.15.0) and two-way ANOVA. Proteins were considered to be regulated by the ABA protocol in a gut microbiota-dependent manner if they showed a significant interaction between gut bacteria depletion and the ABA protocol (*p*(interaction) ≤ 0.05) and met one of the following criteria: (i) differential abundance between CT and ABA mice (|log_2_(fold-change CT vs ABA)| ≥ 0.6), but not between CT-D and ABA-D mice (|log_2_(fold-change CT-D vs ABA-D)| < 0.5), or (ii) differential abundance between CT and ABA mice and opposite differential abundance between CT-D and ABA-D mice ([log_2_(fold-change CT vs ABA) ≥ 0.6 and log_2_(fold-change CT-D vs ABA-D) ≤ −0.6] or ([log_2_(fold-change CT vs ABA) ≤ −0.6 and log_2_(fold-change CT-D vs ABA-D) ≥ 0.6]).

PCA and Z-scored protein intensities from differentially abundant hits were plotted in heatmaps after non-supervised hierarchical clustering using Perseus software. Variable Importance projection (VIP) score were calculated and plotted using the “vip” and “ggplot2” packages in R studio (version 2022.02.2), in order to identify proteins that contributed most to differentiating the ABA and CT groups. Correlations between circulating protein levels and clinical parameters or fecal bacteria relative abundances were determined by computing Spearman rank-based correlations and by adjusting p-values for multiple testing using the Benjamini–Hochberg false discovery rate (FDR) correction.

## Results

Four groups of mice were compared in this study: control mice (CT group), control mice treated with a cocktail of antibiotics in order to deplete their gut microbiota (CT-D group), mice exposed to the Activity-Based Anorexia protocol (ABA group), which consists in a restricted access to food combined with a free access to a running wheel, and ABA mice treated with antibiotics (ABA-D group) (Fig. 1A). Body weight, food intake and physical activity were monitored for each animal. As expected, both ABA and ABA-D groups showed a decrease in body weight and food intake compared to CT and CT-D groups at the end of the protocol (Fig 1B-C). Of note, we observed a significantly lower body weight loss in ABA-D mice compared to ABA mice, as previously observed in other studies^39^. This lower decrease in body weight observed for ABA-D mice is probably explained by their lower physical activity, since no difference was observed in food intake between these groups (Fig. 1D-E).

To verify the effectiveness of gut microbiota depletion triggered by the antibiotics provided in the drinking water, we quantified the total number of Eubacteria in each group at the end of the protocol. As expected, the caecal level of Eubacteria is strongly decreased in CT-D and ABA-D mice compared to CT and ABA, thus confirming the efficient depletion of the gut microbiota by antibiotics (Fig 1F). This depletion is furthermore correlated with an increase in caecal content weight (Fig 1G), as previously observed^40^.

At the end of the protocol, different tissues and organs were collected in order to characterize the proteome remodelling induced in response to the ABA model, and to determine which proteins were regulated in a gut microbiota-dependent manner. For each group of animals, four independent sample pools were randomly generated prior to proteomic analysis for the jejunum, colon and liver samples. Blood samples were analyzed individually.

### Remodelling of the jejunal proteome in response to the ABA model

We first characterized the proteome of jejunal tissues in each group of animals. We identified and quantified a total of 4478 proteins (Fig. S1A). Among them, 1783 proteins exhibit differential abundances between groups (*i.e.* proteins altered in response either to the ABA model [*p*(ABA) ≤ 0.05], or to gut bacteria depletion [*p*(gut microbiota depletion) ≤ 0.05] or both [*p*(interaction) ≤ 0.05]; Table S1). By focusing on the 438 proteins significantly altered in response to the ABA model (*p*(ABA) ≤ 0.05; *i.e.* 10% of quantified proteins)), we identified 19 proteins with a strongly increased abundance (fold-change ≥ 2), and 7 proteins with a strongly decreased abundance (fold-change ≤ 0.5) in the ABA groups compared to the control groups (Fig. 2A, 2C, 2D; Table S2). Functional enrichment analyses indicate that some of these proteins are involved in fatty acids catabolism (Ces1), gluconeogenesis (Pck1 and Fbp1) and adaptive immune responses (Cd74 and H2-Eb1) (Fig. 2E, 2F; Table S3).

**Figure 2:**
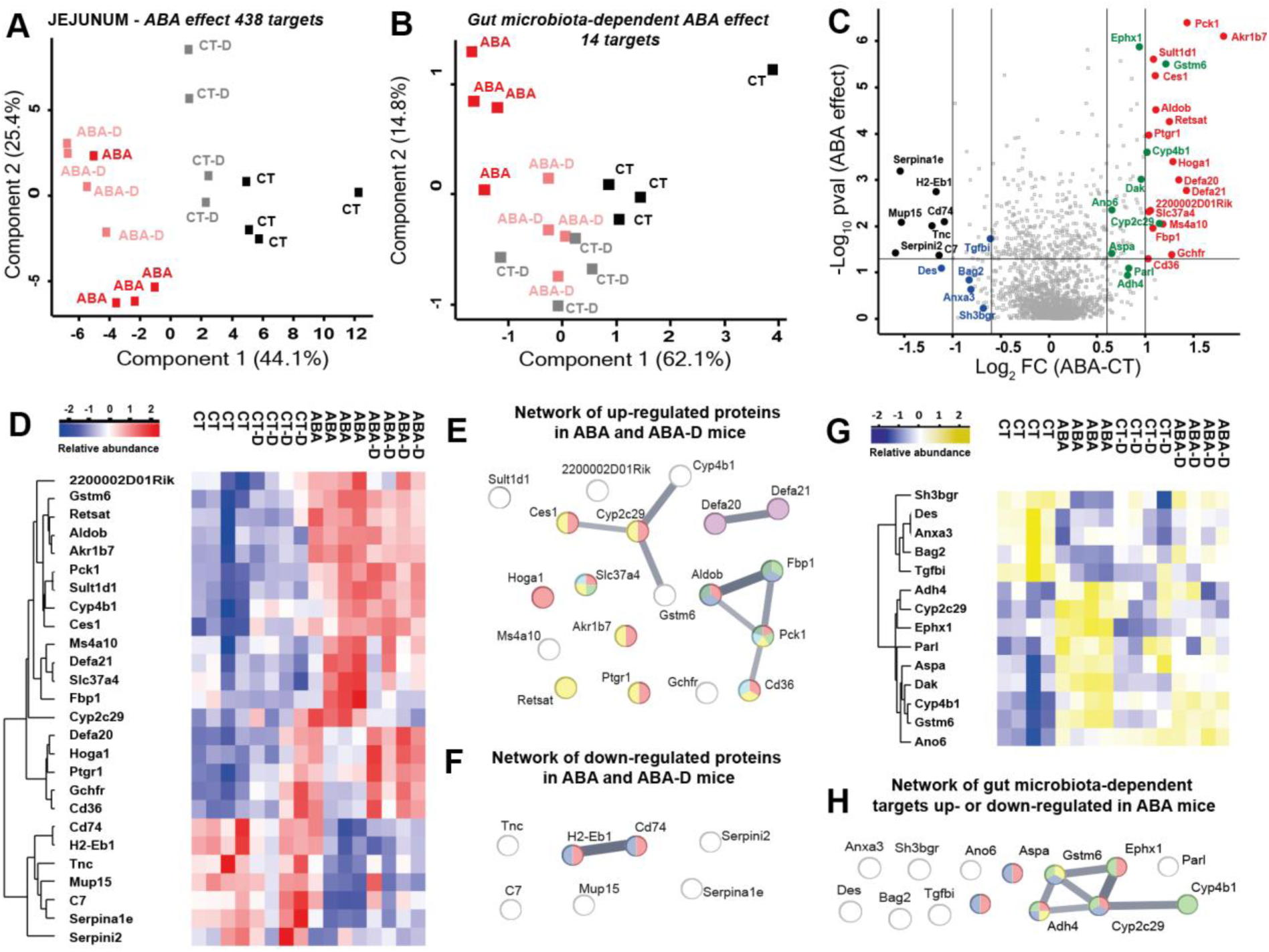
Proteome remodelling in the jejunum triggered in response to the ABA model. (A) Principal component analysis (PCA) based on the 438 jejunal proteins from the jejunum differentially regulated in response to the ABA model (ABA effect). (B) PCA based on the 14 proteins differentially regulated in the ABA group in a gut microbiota-dependant manner (gut microbiota-dependent ABA effect). (C) Volcano plot of the proteins from the jejunum differentially regulated in ABA versus CT groups. Red and black symbols correspond to proteins that are up- and down-regulated, respectively, in the jejunum of ABA mice (ABA effect), while green and blue symbols correspond to proteins up- and down-regulated, respectively, in the jejunum of ABA mice in a gut microbiota-dependant manner (gut microbiota-dependent ABA effect). (D) Heatmap of the top 26 proteins from the jejunum differentially regulated in response to the ABA model. (E) Functional enrichment (FE) analysis highlighting the interactions between proteins up-regulated in response to the ABA protocol (red, monocarboxylic acid metabolic process; blue, fructose 1,6-bisphosphate metabolic process; green, gluconeogenesis; yellow, cellular lipid metabolic process; cyan, triglyceride metabolic process; magenta, defensin/corticostatin family; dark green, glycolysis/gluconeogenesis). (F) FE analysis highlighting the interactions between proteins down-regulated in response to the ABA protocol (dark blue, MHC class II protein complex binding; red, CD4 receptor binding). (G) Heatmap of the 14 proteins from the jejunum regulated in response to the ABA model in a gut microbiota-dependent manner. (H) FE analysis highlighting the interactions between the 14 proteins regulated in ABA mice in a gut microbiota-dependent manner (red, monocarboxylic acid metabolic process; yellow, drug metabolism-cytochrome P450; blue, metabolic pathways; green, biological oxidations).

Then, we focused on jejunal proteins regulated in response to the ABA protocol in a gut microbiota-dependent manner (see Methods section for the selection criteria; Fig. 2B and Table S4). We identified 14 proteins that all exhibit differential abundances between CT and ABA mice, but not between CT-D and ABA-D mice, thereby suggesting an involvement of the gut microbiota in the regulation of these targets. Among these 14 proteins, 5 are down-regulated (log_2_(fold-change) ≤ −0.6) and 9 proteins are up-regulated (log_2_(fold-change) ≥ 0.6) in ABA mice compared to CT mice (Fig. 2C and 2G). Functional enrichment analyses highlights that five of these targets (Ephx1, Tkfc, Cyp2c29, Adh4 and Aspa) are involved in monocarboxylic acid metabolic process and biological oxidations (Fig. 2H and Table S5).

Together, these results show that the ABA model triggers a significant remodelling of the jejunal proteome and that some of these jejunal proteins are modulated in a microbiota-dependent manner.

### Remodelling of the colon proteome in response to the ABA model

We performed similar analyses on colon tissue samples. We observed that food restriction combined with physical activity induces a major proteome remodelling in the colon. Among the 4727 quantified proteins, we identified 3945 proteins exhibiting differential abundances between groups (Fig. S1B and Table S6). By focusing on the 2103 proteins significantly altered in response to the ABA model (*i.e.* 44% of quantified proteins), we identified 19 proteins strongly modulated in response to the ABA model, with 2 proteins up-regulated (fold-change ≥ 2) and 17 proteins down-regulated (fold-change ≤ 0.5) in the ABA groups compared to the control groups (Fig. 3A, 3C, 3D; Table S7). Apart from a small number of proteins related to extracellular signalling, no significantly enriched functional categories were identified in these 19 targets (Figure 3E; Table S8).

**Figure 3:**
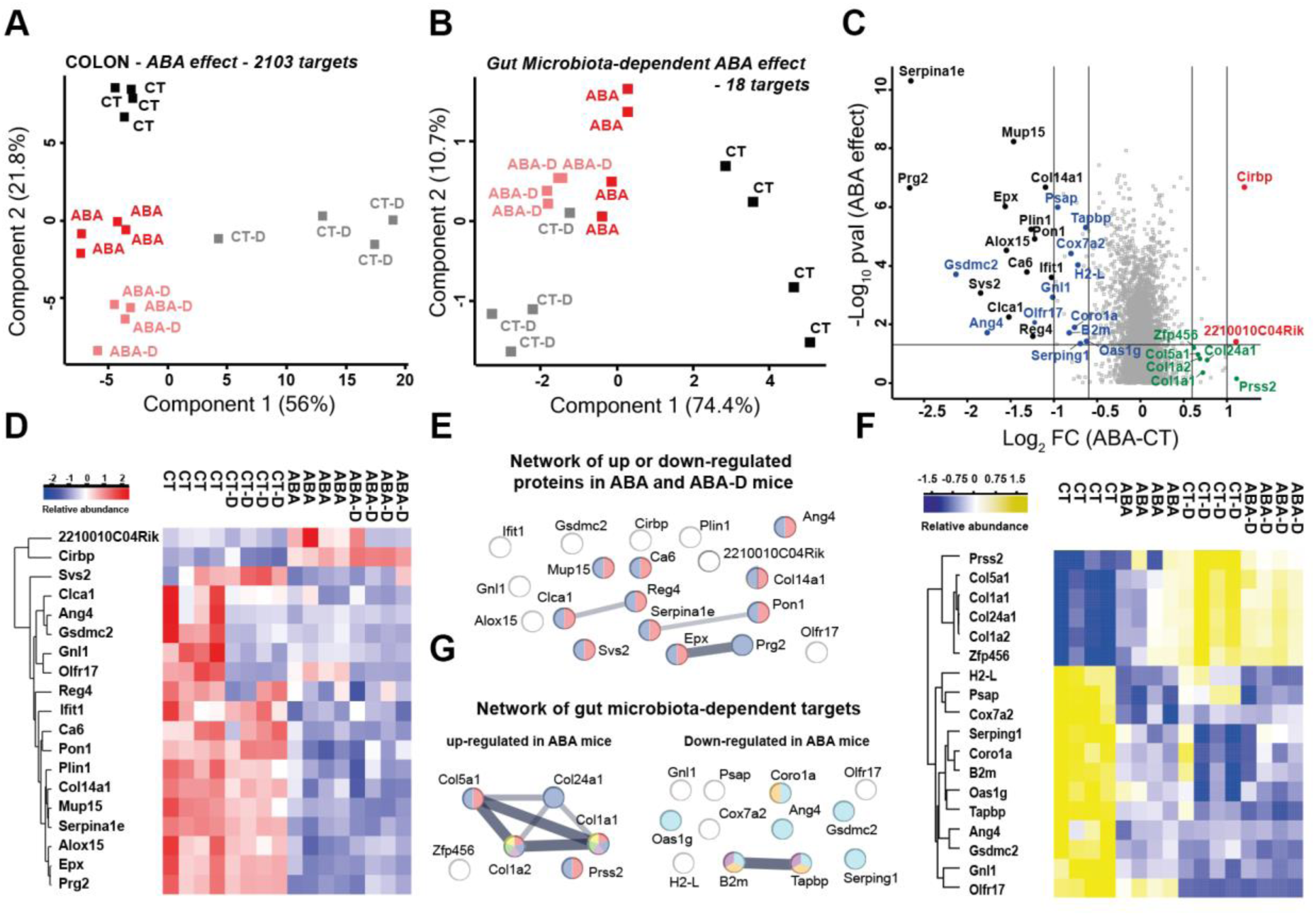
Proteome remodelling in the colon triggered in response to the ABA model. (A) Principal component analysis (PCA) based on the 2103 proteins from the colon differentially regulated in response to the ABA model (ABA effect). (B) PCA based on the 18 proteins differentially regulated in the ABA group in a gut microbiota-dependant manner (gut microbiota-dependent ABA effect). (C) Volcano plot of the proteins from the colon differentially regulated in ABA versus CT groups. Red and black symbols correspond to proteins that are up- and down-regulated, respectively, in the colon of ABA mice (ABA effect), while green and blue symbols correspond to proteins up- and down-regulated, respectively, in the colon of ABA mice in a gut microbiota-dependant manner (gut microbiota-dependent ABA effect). (D) Heatmap of the top 19 proteins from the colon differentially regulated in response to the ABA model. (E) Functional enrichment (FE) highlighting the interactions between proteins up- or down-regulated in response to the ABA protocol (blue, proteins located within the extracellular space; red, proteins acting as a signal molecule). (F) Heatmap of the 18 proteins from the colon regulated in response to the ABA model in a gut microbiota-dependent manner. (G) FE analysis highlighting the interactions between the 18 proteins regulated in ABA mice in a gut microbiota-dependent manner (red, collagen metabolic process; yellow, inflammatory response pathway; blue, protein digestion and absorption; green, dysregulated miRNA targeting in insulin/PI3K-AKT signalling; pink, AGE-RAGE signalling pathway in diabetic complications; cyan, defense response; orange, phagocytic vesicle membrane; purple, MHC class I peptide loading complex).

We then focused on colon proteins regulated in response to the ABA protocol in a gut microbiota-dependent manner, using the same criteria as described above for the jejunum. We identified a total of 18 colon proteins regulated in ABA mice in a gut microbiota-dependent manner (Fig. 3B; Table S9). Among these proteins, 12 are down-regulated in ABA compared to CT mice and mainly involved in immune response mechanisms, and 6 proteins are up-regulated in ABA mice and mainly involved in collagen metabolic process, protein digestion and absorption, and inflammatory responses (Fig 3C, 3F, 3G, Table S10).

Together with the jejunal findings, these results support a role for the gut microbiota in remodeling the colonic proteome in response to the ABA protocol.

### Remodelling of the liver proteome in response to the ABA model

As already observed for the jejunum and the colon, we demonstrated that food restriction combined with physical activity strongly affect the liver proteome. Among the 3823 quantified proteins, we identified 3202 as differentially abundant between groups (Fig. S1C and Table S11). By focusing on the 2562 proteins significantly altered in response to the ABA model (*i.e.* 67% of quantified proteins), we identified 75 liver proteins strongly modulated in response to the ABA protocol, with 24 proteins up-regulated (fold-change ≥ 2) and 51 proteins down-regulated (fold-change ≤ 0.5) in ABA groups compared to control groups (Fig. 4A, 4C, 4D and Table S12). Functional enrichment analyses show that up-regulated proteins in response to the ABA protocol are involved in various pathways including amino acids metabolism, detoxification of copper ion, monooxygenase activity, aryl sulfotransferase activity and PPAR signalling pathways. Liver proteins down-regulated in response to the ABA protocol are involved in fatty acid catabolic metabolism, glucuronosyltransferase activity, epoxygenase P450 pathway and steroid hormone biosynthesis (Fig. 4E, 4F, Table S13).

**Figure 4:**
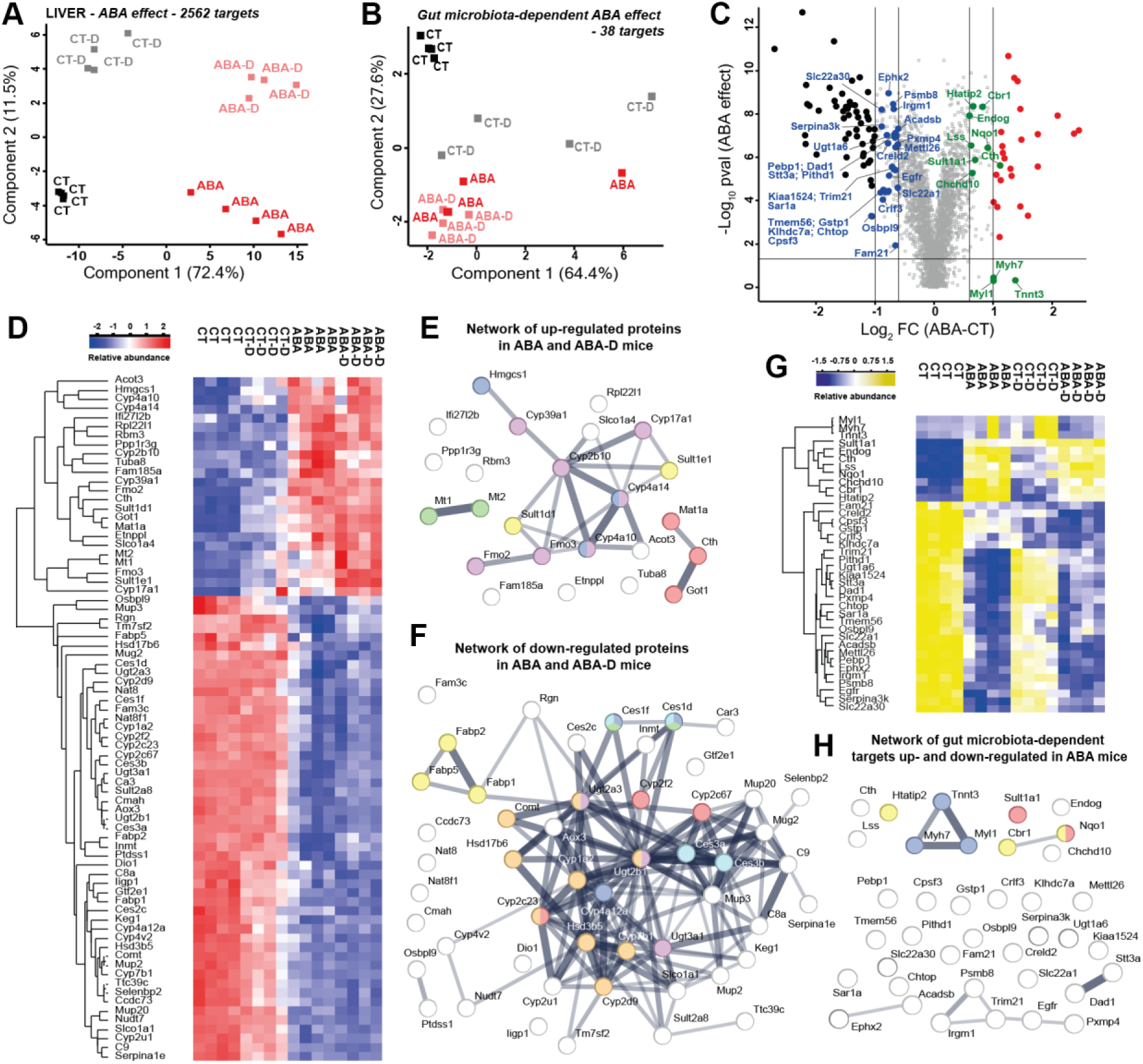
Proteome remodelling in the liver triggered in response to the ABA model. (A) Principal component analysis (PCA) based on the 2562 proteins from the liver differentially regulated in response to the ABA model (ABA effect). (B) PCA based on the 38 proteins differentially regulated in the ABA group in a gut dependant manner (gut microbiota-dependent ABA effect). (C) Volcano plot of the proteins from the liver differentially regulated in ABA versus CT groups. Red and black symbols correspond to proteins that are up- and down-regulated, respectively, in the liver of ABA mice (ABA effect), while green and blue symbols correspond to proteins up- and down-regulated, respectively, in the liver of ABA mice in a gut microbiota-dependant manner (gut microbiota-dependent ABA effect). (D) Heatmap of the top 75 proteins from the liver differentially regulated in response to the ABA model. (E) Functional enrichment (FE) analysis highlighting the interactions between proteins up-regulated in response to the ABA protocol (blue, PPAR signalling pathway; red, biosynthesis of amino acids and cysteine/methionine metabolism; green, detoxification of copper ion; yellow, aryl sulfotransferase activity; magenta, monooxygenase activity). (F) FE analysis highlighting the interactions between proteins down-regulated in response to the ABA protocol (blue, medium-chain fatty acid metabolic process; red, epoxygenase P450 pathway; light blue, methyl indole-3-acetate esterase activity; yellow, long-chain fatty acid transporter activity; magenta, glucuronosyltransferase activity; orange, steroid hormone biosynthesis). (G) Heatmap of the 38 proteins from the liver regulated in response to the ABA model in a gut microbiota-dependent manner. (H) FE analysis highlighting the interactions between the 38 proteins regulated in ABA mice in a gut microbiota-dependent manner (red, estrogen metabolism; blue, muscle protein; yellow, NADP).

Using the same criteria as described above for the jejunum and colon, we found that 38 proteins were regulated in response to the ABA protocol in a gut microbiota-dependent manner (11 up-regulated and 27 down-regulated proteins; Fig. 4B, 4C, 4G and Table S14). Although no functional enrichment was identified among the 27 proteins downregulated in ABA mice in a microbiota-dependent manner, a subset of upregulated proteins are associated with oestrogen metabolism (Nqo1, Sult1a1) and NADP-related energy metabolism (Nqo1, Cbr1 and Htatip2) (Fig. 4H and Table S15).

### Remodelling of the blood proteome in response to the ABA model

We finally determined whether the ABA model triggers a significant remodelling of the blood proteome. Unlike the jejunum, colon and liver analyses, where samples from individual mice were randomly pooled prior to proteomic analysis, blood samples were analyzed here individually. Among the 430 quantified plasma proteins, we identified 278 proteins with differential abundances between groups (Fig. S1D and Table S16). By focusing on the 249 proteins significantly altered in response to the ABA model (*i.e.* 58% of quantified proteins), we identified 136 proteins strongly regulated in response to the ABA model with 124 proteins up-regulated (fold-change ≥ 2) and 12 proteins down-regulated (fold-change ≤ 0.5) in the plasma of ABA groups compared to control groups (Fig. 5A, 5C, S2 and Table S17). Using variable importance projection (VIP) scores, we highlighted the 20 proteins that most strongly discriminate ABA and control animals (Fig. 5C, 5D and 5H).

**Figure 5:**
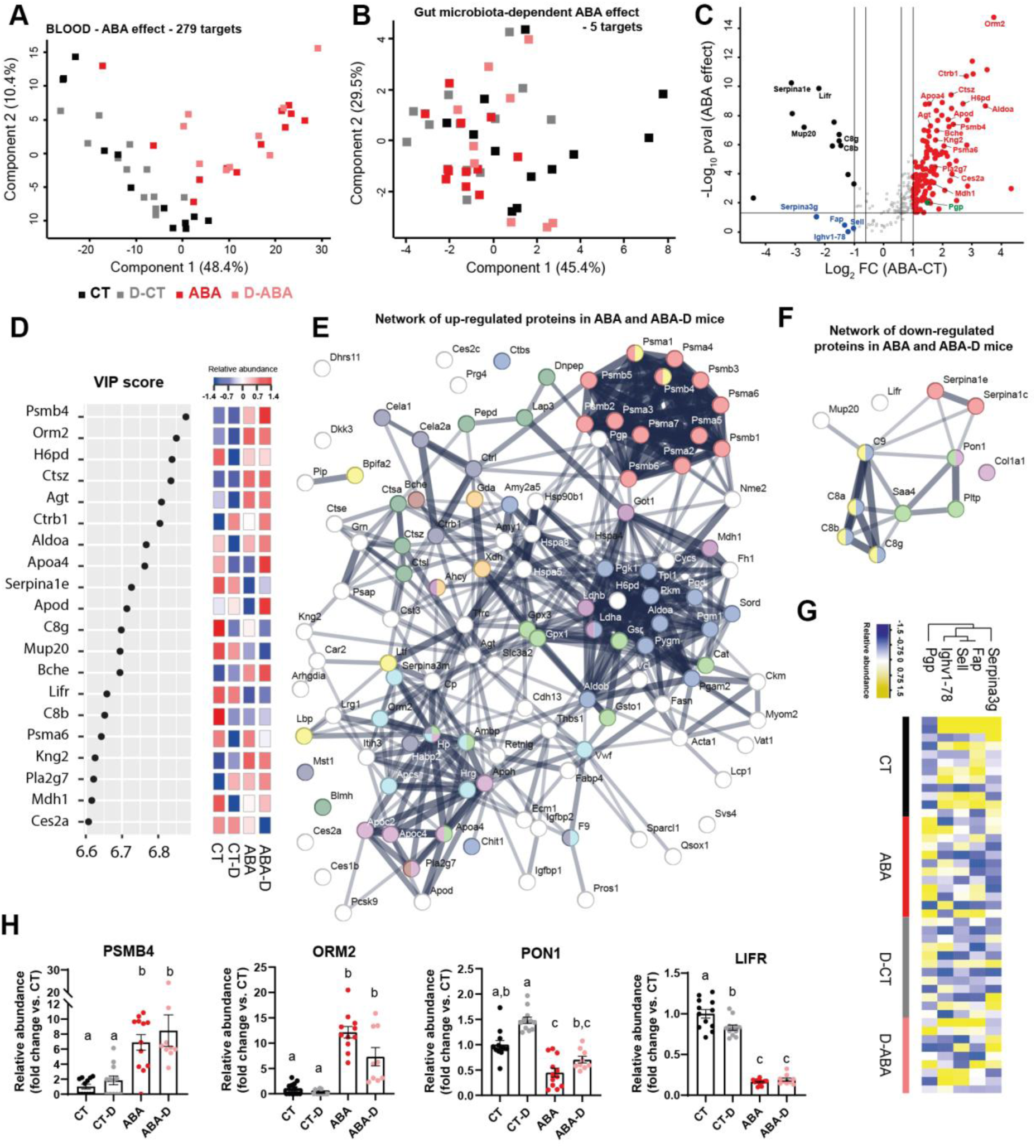
Proteome remodelling in the blood triggered in response to the ABA model. (A) Principal component analysis (PCA) based on the 279 plasma proteins differentially regulated in response to the ABA model (ABA effect). (B) PCA based on the 5 proteins differentially regulated in the ABA group in a gut dependant manner. (C) Volcano plot of the plasma proteins differentially regulated in ABA versus CT groups. Red and black symbols correspond to proteins that are up- and down-regulated, respectively, in the blood of ABA mice (ABA effect), while green and blue symbols correspond to proteins up- and down-regulated, respectively, in the plasma of ABA mice in a gut microbiota-dependant manner (gut microbiota-dependent ABA effect). (D) Variable importance projection (VIP) scores illustrating the top 20 proteins discriminating the ABA and CT groups. (E) Functional enrichment (FE) analysis highlighting the interactions between proteins up-regulated in response to the ABA protocol (cyan, hemostasis, and acute phase; blue, carbohydrate catabolic process; red, proteasome core complex; dark green, exopeptidase activity; light green, antioxidant activity; orange, purine nucleoside catabolic process; yellow, lipopolysaccharide binding; magenta, high-density lipoprotein particle; purple, cysteine and methionine metabolism; dark red, synthesis, secretion, and deacylation of ghrelin; black, trypsin-like serine protease). (F) FE analysis highlighting the interactions between proteins down-regulated in response to the ABA protocol (red, negative regulation of serine-type endopeptidase activity; blue, cytolysis; green, high-density lipoprotein particle; magenta, response to fluoride; yellow, complement activation, classical pathway). (G) Heatmap of the 5 plasma proteins regulated in response to the ABA model in a gut microbiota-dependent manner. (H) Relative abundance of 2 up-regulated proteins (PSMB4 and ORM2) and 2 down-regulated proteins (PON1 and LIFR) in response to the ABA model (PSMB4, ORM2 and PON1, Kruskal-Wallis with Dunn’s multiple comparisons test; LIFR, one-way ANOVA with Tukey’s multiple comparisons test). Different letters indicate statistically significant differences.

Several proteins altered in the plasma of ABA mice are linked in functional networks (Fig. 5E and 5F). Gene ontology and functional enrichment analyses of these proteins are detailed in Table S18 and Fig. S2C. In brief, plasma proteins down-regulated in ABA and ABA-D mice are involved in various biological processes including regulation of endopeptidases, response toward different compounds such as methanol, fluoride and chromate, or High-Density Lipoprotein (HDL) particles metabolism. Interestingly, we found decreased plasma levels of four complement proteins (C8a, C8b, C8g and C9) in ABA mice, suggesting impaired complement-mediated immune function in these animals. Among the 124 proteins up-regulated in the plasma of ABA/ABA-D groups, we identified several proteins involved in the catabolic process of purine nucleoside and carbohydrates, or associated with proteolytic activities (Table S18). We also identified proteins linked to lipopolysaccharide (LPS) binding, antioxidant activity, haemostasis, acute phase response, chylomicron/HDL/very-low-density lipoprotein (VLDL) particles metabolism, and amino acid/pyruvate metabolism. Peptides derived from immunoglobulins (Igha constant domain, and heavy and light chain variable domains) were significantly increased in the plasma of ABA mice. Because immunoglobulin variable regions are not represented in STRING interaction networks, they were excluded from the functional enrichment analysis. Interestingly, we observed that two proteins involved in ghrelin regulation (Bche and Pla2g7) were upregulated in ABA mice blood, suggesting altered signaling of ghrelin, an essential orexigenic hormone, in this model. These two proteins belong to the 20 most discriminant proteins between ABA and control mice according to VIP scores (Fig. 5D).

Finally, we observed a significant increase in the abundance of 13 proteins in the plasma of ABA mice constituting a robust functional network (Psma1-7 and Psmb1-6; Fig. 5E and S2B). These proteins are subunits of the 20S proteasome, which plays a central role in protein turnover and cell stress response. Psam1-7 form the two outer rings of the 20S proteasome, and have essentially structural functions. Psmb1-6 form the two inner rings, which carry the proteasome’s catalytic activity. Our result demonstrates that the ABA model is associated with a significant increase in the level of circulating proteasome.

Using the same criteria as described above for the jejunum, colon and liver, we identified 5 plasma proteins regulated in a gut microbiota-dependent manner in response to the ABA protocol (1 up-regulated and 4 down-regulated in the plasma of ABA mice) (Fig. 5B, 5G, Table S19). No functional enrichment was identified among those proteins.

### Quantification of ORM2 and PON1 plasma levels in patients suffering from Anorexia nervosa

We then aimed to determine whether some of the proteins found to be up- or down-regulated in the blood of ABA mice are also regulated in patients suffering from anorexia nervosa (AN). To this end, we selected two targets displaying opposite regulation in ABA mice, ORM2 (Orosomucoid-2) and PON1 (Serum paraoxonase/arylesterase 1), and quantified their level in plasma samples from patients with AN or from Healthy Volunteers (HV). ORM2, which is an acute-phase protein, is one of the top 20 proteins increased in the blood of ABA mice. PON1, which is a multifunctional protein involved primarily in protection against oxidative stress and inflammation, is in contrast decreased in both plasma and colon in ABA mice. By analysing human blood samples, we show that plasma ORM2 concentration is significantly increased in patients with AN compared to HV (Fig. 6A). In contrast, plasma PON1 concentrations do not differ significantly between the two groups, although a trend toward lower levels is observed in AN patients (p=0.07; Fig. 6B).

**Figure 6:**
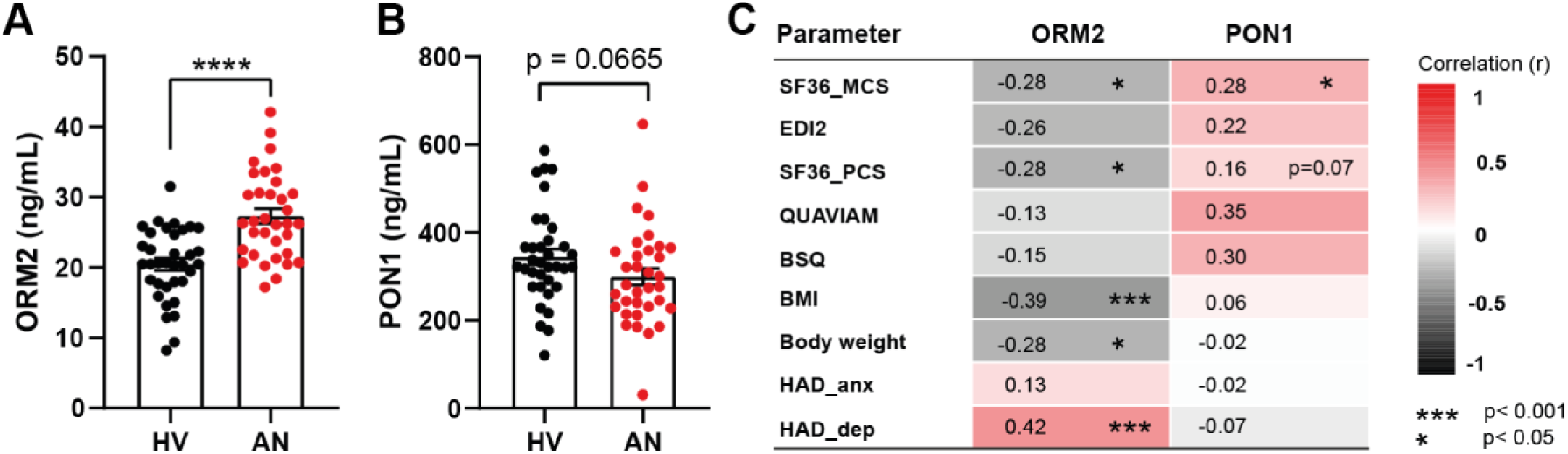
Quantification of ORM2 and PON1 plasma levels of in patients with Anorexia Nervosa and correlations with clinical parameters. (A) Plasma levels of ORM2 in healthy volunteers (HV) and patients with Anorexia Nervosa (AN). (B) Plasma levels of PON1 in HV and patients with AN. Data are presented as individual values with bars representing the mean ± SEM (****, p<0.0001; Mann–Whitney U test). (C) Spearman’s correlation analysis between ORM2 and PON1 plasma levels and clinical parameters, including body mass index (BMI), body weight, Eating Disorder Inventory-2 (EDI-2) questionnaire, Body Shape Questionnaire (BSQ), QUAVIAM questionnaire, Hospital Anxiety and Depression Scale anxiety (HADS-Anx) and depression (HADS-Dep) scores, and SF-36 Physical (PCS) and Mental (MCS) Component Summary scores. Correlation coefficients (r) are represented by a color scale. P-values were adjusted using the Benjamini–Hochberg false discovery rate (FDR) method. *, p<0.05; ***, p<0.001.

We then performed correlation analyses between ORM2 or PON1 levels and individual clinical data, including scores obtained from various questionnaires assessing psychological traits and comorbidities. We observed that plasma level of ORM2 is negatively correlated with individual’s BMI (r = −0.39, p<0.001) and also negatively associated with health-related quality of life (Fig. 6C). Indeed, plasma level of ORM2 is negatively correlated with the Mental Component Score (MCS; r = −0.28, p<0.05) and Physical Component Score (PCS; r = −0.28, p<0.05) from the SF-36 questionnaire^41^, and positively correlated with the Hospital Anxiety and Depression Scale (HADS)/depression score (r = 0.42, p<0.001)^42^ (Fig. 6C). No significant correlations were observed between the plasma level of ORM2 and EDI-2, QUAVIAM, BSQ or HADS/anxiety scores. In the case of PON1, a significant positive correlation was only found with the SF-36 Mental Component Score (r = 0.28, p < 0.05) but not with other clinical data (Fig. 6C).

Finally, we investigated whether circulating levels of ORM2 and PON1 were associated with the relative abundance of specific bacterial taxa in fecal samples from patients and healthy volunteers^15^. No significant correlations were found, indicating that circulating ORM2 and PON1 levels are not driven by the abundance of specific taxa within the gut microbiota. This finding reinforces our observations in mice, where ORM2 and PON1 levels increased in response to the ABA protocol regardless of microbiota status (*i.e.* in both conventional and microbiota-depleted mice; Table S20).

## Discussion

In this study, we used proteomic approaches to explore the changes occurring in the proteome of three organs (*i.e.* jejunum, colon, liver) and one biological matrix (*i.e.* plasma) in response to the ABA model. The rodent ABA model is the most frequently used animal model to mimic AN, as it recapitulates numerous AN features such as low food intake and physical activity, both contributing to a severe body weight loss^43^. We observed in our study a significant decrease in ABA mice body weight, as expected. This decrease is lower in ABA-D mice, which have a depleted gut microbiota, compared to ABA mice, despite similar food intake. This difference has already been observed and has been attributed to a lower physical activity in ABA-D animals^39^. Of note, it has recently been showed that gut bacteria can produce endocannabinoid metabolites that stimulate the release of dopamine in the ventral striatum and increase motivation toward physical activity, contributing *in fine* to negative energy balance and weight loss. Microbiota depletion, in contrast, triggers a decrease in endocannabinoid metabolite levels and is associated with a decrease in exercise capacity^44^. In addition to the body weight loss induced by low food intake, several intestinal dysfunctions have been reported in the ABA model that are also observed in AN patients, including delayed gastric emptying and impairment of gut barrier permeability^31,32,43,45^. A gut dysbiosis has also been described in both ABA mice and anorectic patients, with overlapping features such as an increase in the levels of *Lactobacillus*^13,34^. The ABA model is thus a valuable tool to evaluate anorexia-related proteomic changes and, more particularly, to delineate the role of the gut dysbiosis in these remodelling processes.

Our results showed that food restriction, combined with intensive physical activity, strongly impacts the proteome of various mouse organs and tissues. We observed that proteome remodelling was more pronounced in specific organs of ABA mice, affecting ∼70% of hepatic proteins, ∼60% of plasma proteins and ∼40% of colonic proteins, compared to only 10% of jejunal proteins.

Proteins up-regulated in ABA/ABA-D animals are involved in a wide range of catabolic processes. These results are in accordance with our previous study on the metabolomic changes triggered in ABA mice, where we observed a shift in the metabolism of animals toward muscle breakdown and lower plasma glucose levels compared to control groups^30^. The food restriction imposed in the ABA model triggers body adaptation to catabolic processes promoting the release of glucose and amino acids, thereby ensuring essential physiological functions. Accordingly, previous studies from our group observed higher level of alanine in the plasma and inhibition of protein synthesis in the colon and muscles of ABA mice, associated with an activation of autophagy^30,46,47^. The amino acids released from protein catabolism may be used either for protein resynthesis or for glucose production in other organs or tissues. This is in particular the case for alanine, which may be released from skeletal muscles under long periods of fasting and transferred, via the bloodstream, to the liver, where it allows gluconeogenesis. Of note, we observed an increase in Pck1 level in the jejunum of ABA and ABA-D mice, which may reflect an activation of intestinal gluconeogenesis. Indeed, it has been shown that rats fed with a diet rich in butyrate or in propionate have increased levels of Pck1 in the jejunum, which is associated with the activation of intestinal gluconeogenesis^48^. Our proteomic analyses suggest that lipid metabolism is also altered in ABA/ABA-D animals (in the jejunum and the liver). Altered lipid metabolism has already been observed in the colon of ABA mice as well as in patients with AN^47,49^. The observed shift in metabolism toward catabolic processes may in addition lead to an overaccumulation of oxygen reactive species and ultimately triggers the onset of severe cellular stress, as already observed for example in the stomach of ABA mice^31^. Of note, the PON1 protein, which is downregulated in both ABA/ABA-D plasma and colon, is known to exert both antioxidant and anti-inflammatory activities^50^. Altogether, these data suggest a pro-oxidant state in response to the ABA model.

Regarding the control of eating behaviour, we observed that 2 proteins, Bche (Butyrylcholinesterase) and Pla2g7 (Platelet-activating factor aceytlhydrolase) are up-regulated in the blood of ABA/ABA-D mice. These two enzymes regulate the activity of ghrelin, an orexigenic peptide that requires acylation to bind its receptor. We can hypothesize that the observed increase in Bche and Pla2g7 in ABA/ABA-D mice leads to an increase in the ratio of blood des-acyl/acyl-ghrelin, which may participate to the alteration of eating behaviour observed in this animal model of anorexia^51^. This hypothesis is in line with our previous study, in which we demonstrated an increase in the plasma des-acyl/acyl-ghrelin ratio in ABA mice compared with *ad libitum*-fed mice^52^.

Interestingly, several proteins involved in immune responses were observed as dysregulated in ABA mice. 9 of the up-regulated plasma proteins in ABA mice (Vwf; F9; Hrg; Serpina3m; Ambp; Apcs; ORM2; Hp; Itih3) belong to the acute phase and coagulation responses, including serum amyloid P component (Apcs), the functional counterpart of human C-reactive protein. We also observed an up-regulation of the resistin like γ (Retnlg) protein in the plasma of ABA mice, which is also over-expressed in a wide range of diseases including inflammatory diseases and metabolic disorders^53^. These observations are in line with the previously reported low-grade inflammatory state observed in AN. Additionally, we observed the downregulation of several other immune-related proteins, including Prg2, Epx, Alox15, Reg4, Ifit1, Clca1 and PON1, that likely reflects a modulation in immune responses. This is consistent with the well-documented impact of undernutrition on immune functions. Indeed, limited energy availability leads to a reprioritization of physiological processes, while the modulation of peptides such as leptin may directly impact immune cell activity by regulating glucose utilization^54^. The downregulation of the complement C8 and C9 proteins in the blood further points to a potential impairment of pathogen defense in ABA animals. This impairment of immune response capacities should be considered in light of the alteration of the intestinal barrier in these animals. Indeed, an increase in intestinal barrier permeability has been observed in ABA mice, which may promote the translocation of bacterial components or opportunistic pathogens from the intestinal lumen to deeper tissues and elicit inflammatory responses^32^.

One intriguing finding of our proteomic analyses in blood samples is the sharp increase in the level of circulating 20S proteasome subunits. The role of circulating proteasome subunits is poorly characterized. Increased levels of circulating 20S proteasome subunits have been observed in mouse under pathophysiological conditions and in patients suffering from several diseases including auto-immune disorders, blood, skin and solid cancers, trauma and sepsis^55,56^. However, the origin and the biological roles or mechanisms of extracellular release of these proteasomal subunits still remain elusive^55^. The potential causes and consequences of this increase in circulating 20S proteasome in AN deserve further investigations.

A key finding of our study is that a significant number of proteins are regulated in ABA mice in a gut microbiota-dependent manner. Indeed, we identified 14, 18, 38 and 5 proteins in the jejunum, colon, liver and blood, respectively, that were differentially altered in ABA mice and ABA-D mice. These results suggest that these proteins are regulated either by signals emitted by the dysbiotic microbiota associated with the ABA model, or directly by undernutrition and physical activity acting in synergy with gut microbiota-derived signals. Notably, we cannot rule out that the differential regulation of certain proteins is indirectly linked to the reduced body weight loss observed in microbiota-depleted ABA mice.

In the jejunum, five of the proteins up-regulated in ABA mice in a microbiota-dependent manner are involved in lipid and monocarboxylic acid metabolism (Tkfc, Ephx1, Cyp2c29, Adh4 and Aspa). A growing body of evidence has demonstrated that gut bacteria play a pivotal role in modulating host lipid homeostasis^57–59^. Two of these proteins act at consecutive steps of a single pathway: Cyp2c29 is an arachidonate 14,15-epoxygenase that generates fatty acid epoxides such as epoxyeicosatrienoic acids, and microsomal epoxide hydrolase (Ephx1) hydrolyses these epoxides to their corresponding diols, contributing substantially to their turnover *in vivo*^60,61^. This is the case for example of the lipid-derived mediator arachidonic acid, which can be metabolized by cytochrome P450 epoxygenases and subsequently converted into dihydroxyeicosatrienoic acids by epoxide hydrolases. Because increased epoxygenase and increased epoxide hydrolase activities act in opposite directions, the net consequence on the jejunal epoxide:diol ratio cannot be inferred from protein abundance alone.

In the colon, our results point to a microbiota-dependent remodelling of the mucosal barrier and immune functions. Collagens I and V (Col1a1, Col1a2, Col5a1) were increased in ABA mice, indicating extracellular matrix remodelling. We can hypothesize that this increase reflects *de novo* fibrillogenesis, a hypothesis supported by the increase in collagen V, which is a quantitatively minor but regulatory collagen required for the nucleation of collagen I fibrils. This mechanism is likely gut microbiota-dependent since increases in collagen proteins are observed in ABA mice and not in ABA-D mice. Of note, collagen deposition has been described in intestinal injury models in which dysbiosis contributes to low-grade inflammation and tissue remodelling^62^. The downregulation of antimicrobial factors, such as Ang4, and proteins involved in antigen presentation (B2m, Tapbp, H2-L) in ABA mice suggest in addition a decrease in mucosal immune surveillance capacities in these animals, triggered by microbiota-dependent factors. Ang4 is a *bona fide* microbiota-inducible effector as its expression can be modulated by *Bacteroides* bacteria^63^. Decreases in Gnl1 and Ang4 levels could thus reflect a gut microbiota-dependent modulation of epithelial renewal in the ABA model^64–66^. B2M, Tapbp and MHC class I heavy chains are co-regulated by IFNγ. Their coordinated decrease could be interpreted as a decrease in the IFNγ response at the level of the colonic epithelium.

In the liver, we identified 38 proteins regulated in a microbiota-dependent manner. Several up-regulated proteins in ABA mice converge on xenobiotic detoxification and redox control such as the phase-II sulfotransferase Sult1a1, the quinone reductase Nqo1 and carbonyl reductase 1 (Cbr1). Nqo1 and Cbr1 are both established NRF2 targets^67,68^. Their coordinated increase suggests a microbiota-dependent activation of the NRF2 antioxidant programme. This is consistent with the previously established regulation of hepatic phase-II enzyme expression by the microbiota, which is markedly altered in germ-free animals and restored by conventionalisation^69^. In addition, we established that Endonuclease G (Endog), which is a positive regulator of starvation-induced autophagy, is up-regulated in the liver of ABA mice only. This observation fits with the previously reported liver damage associated with autophagy reported in ABA mice and patients with AN^70,71^. While starvation-induced autophagy has previously been implicated in liver alterations associated with AN, our proteomic data additionally revealed changes in proteins involved in apoptosis-related pathways, including decreases in Dad1 and increases in Htatip2 and ChChd10^72,73–75^. Overall, these findings suggest a model in which microbiota-dependent signals reach the liver and modulate hepatic redox defence, autophagy and apoptosis-related pathways in response to chronic food restriction.

We identified five proteins (Serpina3g, Fap, Sell, Ighv1-78, and Pgp) that were specifically modulated in the plasma of ABA mice and not in ABA-D mice. Although no significant functional enrichment was detected, several of these proteins appear to be linked to immune-related processes. For instance, L-selectin (encoded by the *sell* gene) plays a pivotal role in immune responses. Its soluble form, generated by the cleavage of membrane-bound L-selectin, can modulate leukocyte-endothelial interactions by competing with membrane-bound L-selectin for its ligands, thereby limiting leukocyte rolling and adhesion^76^. In contrast to such specialized mediators, many of our identified targets are primarily cytosolic or membrane-anchored, suggesting that their altered plasma abundance may reflect changes in tissue turnover or cellular release rather than a direct systemic function. Consequently, determining the functional implications of these circulating fluctuations remains challenging.

To assess the translational relevance of our preclinical findings, we quantified the serum levels of two proteins, ORM2 and PON1, which were identified as differentially abundant in our ABA mouse model, in a subset of patients with Anorexia. ORM2 encodes an acute-phase protein that is upregulated during inflammatory responses. Its immunological effect is context-dependent as it may increase cytokine-mediated inflammation in rheumatoid synovium, or have anti-inflammatory effects in models of steatohepatitis^77,78^. We demonstrate here that ORM2 constitutes an interesting biomarker in the context of anorexia. Indeed, this protein displays significantly higher levels in the serum of AN patients compared to healthy volunteers. Moreover, we show that increased ORM2 levels correlate with lower BMI and body weight. Interestingly, it has been demonstrated that ORM2 deficiency exacerbates high-fat diet-induced obesity and intestinal inflammation in mice, thereby strengthening our correlation between ORM2 levels and BMI^79^. We also observed that increased ORM2 levels correlate with poorer scores on physical and mental health-related questionnaires (SF36 and HADS). Together, these findings suggest that ORM2 levels reflect the severity of malnutrition and are associated with some of the psychological traits characteristic of Anorexia Nervosa. In contrast, circulating PON1 concentrations did not differ significantly between AN patients and healthy volunteers, although they showed a trend toward lower levels in patients. Interestingly, we observed that higher PON1 concentrations were associated with better mental health-related quality of life. However, given the absence of a significant difference between patients and controls, these observations should be interpreted cautiously and warrant confirmation in larger cohorts. No significant associations were observed between circulating ORM2 or PON1 concentrations and the relative abundance of bacterial taxa in patients healthy volunteers’ fecal samples. These findings are consistent with our experimental data demonstrating that both proteins are modulated in a microbiota-independent manner in ABA mice and probably belong to the animal’s response to undernutrition.

Our proteomic results should be considered with some limitations. Conducting an analysis at the end of the ABA protocol prevents us from accessing the dynamics of proteome remodelling in response to food restriction and physical activity. In addition, our experimental approach is based on the analysis of whole tissues which prevents the identification of cell-type specific protein regulation. Delineating the sex-dependent differences in organ proteome remodelling would also be interesting since sex-dependent responses were observed in the ABA model^80,81^.

Altogether, our results demonstrate that food restriction and physical activity, two hallmarks of Anorexia Nervosa, have a strong impact on the proteome of various organs and tissues. Our study provides a valuable dataset for advancing our understanding of the pathophysiological mechanisms underlying this disease. Furthermore, some of the targets identified in the bloodstream may constitute promising biomarkers to help predict either chronicization or remission of the disease. Our study further highlights the role of the gut microbiota in modulating specific proteins across different organs, thereby providing new insights into its potential contribution to the pathophysiology of Anorexia Nervosa.

## Supporting information

Graphical Abstract

## Ethics approval

The study design has been approved by the French national committee “Informatique et liberté” (N$ CNIL: 1787487) and the Ethics Committee (N°CPP/CE 002/2014). Patients and healthy individuals agreeing to participate provided a written informed consent.

## Data sharing

The mass spectrometry proteomics data have been deposited to the ProteomeXchange Consortium via the PRIDE partner repository under the dataset identifiers PXD086112 (TMT-based quantitative proteomics of three organ types) and PXD086168 (label-free quantitative plasma proteomics).

## Acknowledgements

We thank the VIB Proteomics Core for help with the proteome analysis.

## Figure legends (supplementary data)

***Supplemental Figure S1: Principal Component Analysis (PCA) based on proteomic data from different tissues.*** PCA plots based on all quantified proteins from jejunum (A), colon (B), liver (C) and plasma (D) samples. Black, CT group; grey, CT-D group; red, ABA group; pink, ABA-D group.

***Supplemental Figure S2: Remodelling of the blood proteome in response to the ABA protocol.*** (A) Heatmap of the top 136 blood proteins differentially regulated in response to the ABA model. (B) Heatmap illustrating the abundance of Psma and Psmb proteasome subunits in individual animals from the CT, CT-D, ABA and ABA-D groups. (C) Main functional annotations of the blood proteins regulated in response to the ABA model.

