## Supplemental Figure S1 for "Role of the gut microbiota in the remodelling of the jejunal, colonic, hepatic and blood proteomes in a mouse model of Anorexia Nervosa"

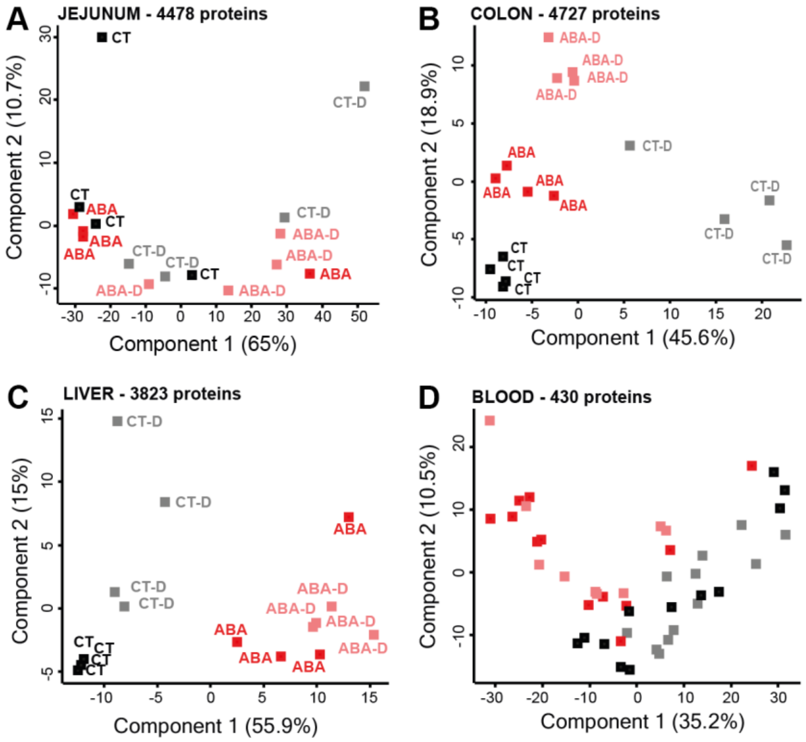
A
JEJUNUM - 4478 proteins
Component 2 (10.7%)
Component 1 (65%)
B
COLON - 4727 proteins
Component 2 (18.9%)
Component 1 (45.6%)
C
LIVER - 3823 proteins
Component 2 (15%)
Component 1 (55.9%)
D
BLOOD - 430 proteins
Component 2 (10.5%)
Component 1 (35.2%)
CT
CT-D
ABA
ABA-D

### Figure S1
