## Supplemental Figure S2 for "Role of the gut microbiota in the remodelling of the jejunal, colonic, hepatic and blood proteomes in a mouse model of Anorexia Nervosa"

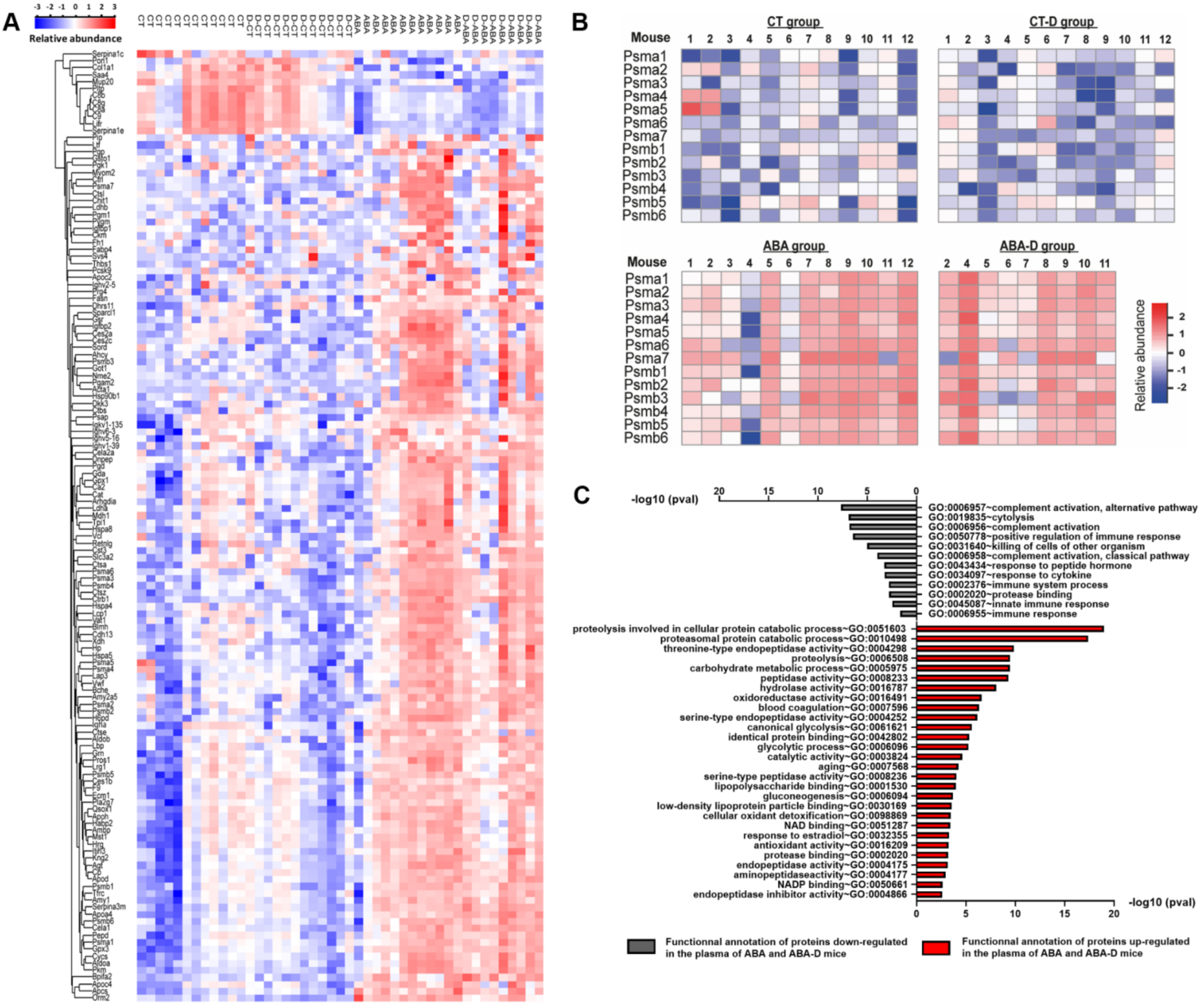
A
Relative abundance
B
CT group
CT-D group
ABA group
ABA-D group
Mouse
Psma1
Psma2
Psma3
Psma4
Psma5
Psma6
Psma7
Psmb1
Psmb2
Psmb3
Psmb4
Psmb5
Psmb6
Relative abundance
C
-log10 (pval)
GO:0006957~complement activation, alternative pathway
GO:0019835~cytolysis
GO:0006956~complement activation
GO:0050778~positive regulation of immune response
GO:0031640~killing of cells of other organism
GO:0006958~complement activation, classical pathway
GO:0043434~response to peptide hormone
GO:0034097~response to cytokine
GO:0002376~immune system process
GO:0002020~protease binding
GO:0045087~innate immune response
GO:0006955~immune response
proteolysis involved in cellular protein catabolic process~GO:0051603
proteasomal protein catabolic process~GO:0010498
threonine-type endopeptidase activity~GO:0004298
proteolysis~GO:0006508
carbohydrate metabolic process~GO:0005975
peptidase activity~GO:0008233
hydrolase activity~GO:0016787
oxidoreductase activity~GO:0016491
blood coagulation~GO:0007596
serine-type endopeptidase activity~GO:0004252
canonical glycolysis~GO:0061621
identical protein binding~GO:0042802
glycolytic process~GO:0006096
catalytic activity~GO:0003824
aging~GO:0007568
serine-type peptidase activity~GO:0008236
lipopolysaccharide binding~GO:0001530
gluconeogenesis~GO:0006094
low-density lipoprotein particle binding~GO:0030169
cellular oxidant detoxification~GO:0098869
NAD binding~GO:0051287
response to estradiol~GO:0032355
antioxidant activity~GO:0016209
protease binding~GO:0002020
endopeptidase activity~GO:0004175
aminopeptidaseactivity~GO:0004177
NADP binding~GO:0050661
endopeptidase inhibitor activity~GO:0004866
-log10 (pval)
Functionnal annotation of proteins down-regulated in the plasma of ABA and ABA-D mice
Functionnal annotation of proteins up-regulated in the plasma of ABA and ABA-D mice

**Figure S2**
