## Supplemental Methods for "Role of the gut microbiota in the remodelling of the jejunal, colonic, hepatic and blood proteomes in a mouse model of Anorexia Nervosa"

[1] Univ Rouen Normandie, INSERM, Normandie Univ, ADEN, UMR 1073 "Nutrition, Inflammation and Microbiota-Gut-Brain axis", Rouen, France.

[2] CHU Rouen, Department of Nutrition, Rouen, France

[3] Department of Biomolecular Medicine, Ghent University, Ghent, Belgium.

[4] VIB Center for Medical Biotechnology, Ghent, Belgium.

[5] VIB Proteomics Core, Ghent, Belgium.

[6] Univ Rouen Normandie, INSERM US51, CNRS UAR2026, Behavioural Analysis Platform SCAC-HeRacLeS, Rouen, France.

[7] CHU Rouen, CIC-CRB1404, Rouen, France.

[8] CHU Rouen, Department of Biochemistry, Rouen, France

*Corresponding authors:
Jonathan Breton - INSERM U1073, UFR Santé, 22 boulevard Gambetta, 76183 ROUEN cedex France –
David Ribet - INSERM U1073, UFR Santé, 22 boulevard Gambetta, 76183 ROUEN cedex France –

*Sample preparation for proteomics analyses*

Jejunal, colonic and hepatic samples were lysed in 1 ml lysis buffer (5% sodium dodecyl sulfate (SDS), 50 mM triethylammonium bicarbonate (TEAB), pH 8.5) in gentleMACS M tubes. Lysis was performed with a gentleMACS® dissociator and the lysed tissue was centrifuged for 5 min at 4,000 x g. Next, samples were sonicated with 3 pulses of 15 s at an amplitude of 20% using a 3 mm probe, with incubation on ice for 1 minute between pulses. After centrifugation for 15 minutes at 7,197 x g at room temperature (RT), to remove insoluble components, the protein concentration was measured by bicinchoninic acid (BCA) assay (Thermo Scientific). In each group of animals, and for each type of segments, we pooled three to four samples from independent animals randomly[1]. 100 µg of protein from each pooled sample were reduced by adding 15 mM dithiothreitol (30 minutes at 55˚C) and then alkylated by adding 30 mM iodoacetamide (15 minutes at RT in the dark). Phosphoric acid was then added to a final concentration of 1.2%. Samples were diluted 7-fold in a binding buffer containing 90% methanol in 100 mM TEAB, pH 7.55. The samples were loaded on a 96-well S-Trap plate (Protifi), placed on top of a deepwell plate, and centrifuged for 2 min at 1,500 x g at RT. After protein binding, the S-trap plates were washed three times with binding buffer and centrifugated for 2 min at 1,500 x g at RT. A new deepwell receiver plate was placed below the 96-well S-Trap plate and 50 mM TEAB containing trypsin (1/100, w/w) was added for digestion overnight at 37°C. Three rounds of elution were finally performed to elute peptides, first with 80 µl 50 mM TEAB, then with 80 µl 0.2% formic acid in water and finally with 80 µl 0.2% formic acid in water/acetonitrile (ACN) (50/50, v/v). Eluted peptides were dried completely by vacuum centrifugation.

Peptides obtained from jejunal, colonic and hepatic segments were further TMT (Tandem Mas Tag)-labelled. TMTpro 16-plex labels (0.5 mg, Thermo Fisher Scientific) were equilibrated to RT immediately before use and dissolved in 25 µl anhydrous ACN. Dried peptides were re-suspended in 20 µl 100 mM TEAB (pH 8.5), peptide concentration was determined on a Lunatic spectrophotometer (Unchained Labs)[2] and 30 µg (colon/jejunum) or 50 µg (liver) were labelled for 1 hour at RT using 0.1 mg (5 µl) of TMTPro label (labels used: 126C, 127N, 127C, 128N: CT replicates; 128C, 129N, 129C, 130N: ABA replicates; 130C, 131N, 131C, 132N: D-CT replicates; 132C, 133N, 133C, 134N: D-ABA replicates). The reaction was quenched for 15 min at RT by addition of 1 µl 5% hydroxylamine. The 16 labelled samples were combined and 100 µg of mixed labelled peptides were dried by vacuum centrifugation, re-dissolved in 200 µl solvent A (0.1% TFA in water/ACN (98:2, v/v)) and desalted on a reversed phase (RP) C18 OMIX tip (Agilent). The tip was first washed 3 times with 200 µl pre-wash buffer (0.1% TFA in water/ACN (20:80, v/v)) and pre-equilibrated 5 times with 200 µl of wash buffer (0.1% TFA in water) before the sample was loaded on the tip. After peptide binding, the tip was washed 3 times with 200 µl of wash buffer and peptides were eluted twice with 100 µl elution buffer (0.1% TFA water/ACN (40:60, v/v)). The combined elutions were dried in a vacuum concentrator. Vacuum dried peptides were re-dissolved in 100 µl solvent A and 95 µl was injected for fractionation by RP-HPLC (Agilent series 1200) connected to a Probot fractionator (LC Packings). Peptides were first loaded in solvent A on a 4 cm pre-column (made in-house, 250 µm internal diameter (ID), 5 µm C18 beads, Dr. Maisch) for 10 min at 25 µl/min and then separated on a 15 cm analytical column (made in-house, 250 µm ID, 3 µm C18 beads, Dr Maisch). Elution was done using a linear gradient from 100% RP-HPLC solvent A (10 mM ammonium acetate (pH 5.5) in water/ACN (98:2, v/v)) to 100% RP-HPLC solvent B (70% ACN, 10 mM ammonium acetate (pH 5.5)) in 100 min at a constant flow rate of 3 µL/min. Fractions were collected every minute between 20 and 92 minutes and pooled every 12 minutes to generate a total of 12 samples for LC-MS/MS analysis. All 12 fractions were dried under vacuum in HPLC inserts and stored at −20°C until further use.

For plasma samples, 1 µl of samples was mixed with 50 µl of lysis buffer containing 5% SDS and 50 mM TEAB, pH 8.55 and processed with the S-trap protocol as described above.

*LC-MS/MS analyses*

TMT-labelled peptides from jejunal, colonic and hepatic segments were re-dissolved in 20 µl solvent A of which 15 µl was injected for LC-MS/MS analysis on an Ultimate 3000 RSLCnano system (Thermo) in line connected to a Q Exactive HF Biopharma mass spectrometer (Thermo) equipped with a pneu-Nimbus dual ion source (Phoenix S&T). Trapping was performed at 10 µl/min for 4 min in solvent A on a 20 mm trapping column (made in-house, 100 µm internal diameter (I.D.), 5 µm beads, C18 Reprosil-HD, Dr. Maisch, Germany). The peptides were separated on a 250 mm nanoEase column (M/Z HSS T3 column, 100 Å, 1.8 µm, 75 µm I.D., Waters) kept at a constant temperature of 40°C. Peptides were eluted by a non-linear increase from 1 to 55% MS solvent B (0.1% formic acid (FA) in water/ACN (2:8, v/v)) over 105 minutes, at a flow rate of 300 nl/min, followed by a 15-minutes wash reaching 97% MS solvent B and re-equilibration with 99% MS solvent A (0.1% FA in water). The mass spectrometer was operated in data-dependent mode, automatically switching between MS and MS/MS acquisition for the 15 most abundant ion peaks per MS spectrum. Full-scan MS spectra (375-1,500 m/z) were acquired at a resolution of 120,000 in the Orbitrap analyzer after accumulation to a target value of 3,000,000. The 15 most intense ions above a threshold value of 10,000 (minimum AGC of 1,000) were isolated for fragmentation at a normalized collision energy of 35%. The C-trap was filled at a target value of 100,000 for maximum 100 ms and the MS/MS spectra (200-2,000 m/z) were acquired at a resolution of 45,000 in the Orbitrap analyzer with a fixed first mass of 110 m/z. Only peptides with charge states ranging from +2 to +6 were included for fragmentation and the dynamic exclusion was set to 12 s. QCloud was used to control instrument longitudinal performance during the project[3].

Peptides from plasma samples were run in data-independent mode on an Evosep One LC-system (Evosep, Denmark) in-line connected to a Q Exactive HF mass spectrometer (Thermo). Peptides were analyzed with the 30 SPD method using the endurance Evosep column (15cm x 150 µm I.D., 1.9 µm beads, Evosep, Denmark). For elution of the peptides from the column 0.1% FA in LC-MS-grade water can 0.1% FA in ACN were used as mobile phases.

Full-scan MS spectra ranging from 375-1500 *m/z* with an AGC target value of 5,000,000, a maximum fill time of 50 ms and a resolution at 200 m/z of 60,000 were followed by 30 quadrupole isolations with a precursor isolation width of 10 *m/z* for HCD fragmentation at an NCE of 30% after filling the trap at a target value of 3,000,000 for maximum injection time of 45 ms. MS2 spectra were acquired at a resolution of 15,000 at 200 *m/z* in the Orbitrap analyser without multiplexing. The isolation intervals ranging from 400 – 900 *m*/*z*, with an overlap of 5 *m*/z were created with the Skyline software tool. QCloud was used to control instrument longitudinal performance during the project[3].

*Proteomics data analysis*

Data analysis of the tissue samples was performed in MaxQuant (version 1.6.2.10) with mainly default search settings including a false discovery rate set at 1% on PSM (peptide-spectrum match) and protein level. Spectra were searched against the mouse reference proteome (database release version of 2021_01, containing 21,989 sequences, www.uniprot.org). The mass tolerance for precursor and fragment ions was set to 4.5 and 20 ppm, respectively, during the main search. Enzyme specificity was set as C-terminal to arginine and lysine, also allowing cleavage at proline bonds with a maximum of two missed cleavages. Variable modifications were set to oxidation of methionine residues and acetylation of protein N-termini whereas carbamidomethylation of cysteine residues was set as fixed modification. In order to reduce the number of spectra that suffer from co-fragmentation, the precursor ion fraction (PIF) option was set to 75%. Only proteins with at least one unique or razor peptide were retained. MS2-based quantification using TMTpro labels was chosen as quantification method and a minimum ratio count of two unique peptides was required for quantification. Further data analysis was performed with an in-house R script. Reverse database hits were removed from the MaxQuant protein groups file, corrected reporter intensities were $\log_2$ transformed & median normalized and replicate samples were grouped. Proteins with less than three valid values in at least one group were removed and missing values were imputed by random sampling from a normal distribution centered around each sample's detection limit (package DEP)[4] leading to a list of 4478, 4727 and 3823 quantified proteins in jejunum, colon and liver samples respectively, that were used for further data analysis.

For data analysis of the plasma samples, LC-MS/MS runs of all 45 samples were searched together using the DiaNN algorithm (version 1.8) with mainly default search settings, including a false discovery rate set at 1% on precursor peptide and protein level. Spectra were searched against the mouse protein sequences in the Swiss-Prot database (database release version of 2021_01), containing

21,989 sequences (www.uniprot.org). Carbamidomethylation of cysteines, oxidation of methionine residues and acetylation of the protein Ntermini were set as variable through the additional options. Trypsin was used as the digestion enzyme, allowing cleavage N-terminally at proline and allow for two missed cleavages. Mainly default settings were used, except for the addition of a 375-900 m/z precursor mass range filter, the match between runs (MBR) option and MS2 mass accuracy was set to 20 ppm and MS1 mass accuracy to 10 ppm. Further data analysis was performed with an in-house script in the R programming language. The DIA-NN main report output table was filtered at a precursor and protein library q-value cut-off of 1 % and only proteins identified by at least one proteotypic peptide were retained. MaxLFQ intensities were $log_2$ transformed and replicate samples were grouped. Proteins with less than three valid values in at least one group were removed and missing values were imputed from a normal distribution centered around the detection limit (package DEP)[4] leading to a list of 430 quantified proteins in the experiment, used for further data analysis.

STRING (version 12.0) and DAVID (DAVID 2021, DAVID Knowledgebase: v2023q4) were used as bioinformatic resources for functional annotation and enrichment analyses of the identified proteins.
