## Supplementary material for "Role of the gut microbiota in the remodelling of the jejunal, colonic, hepatic and blood proteomes in a mouse model of Anorexia Nervosa": Graphical Abstract

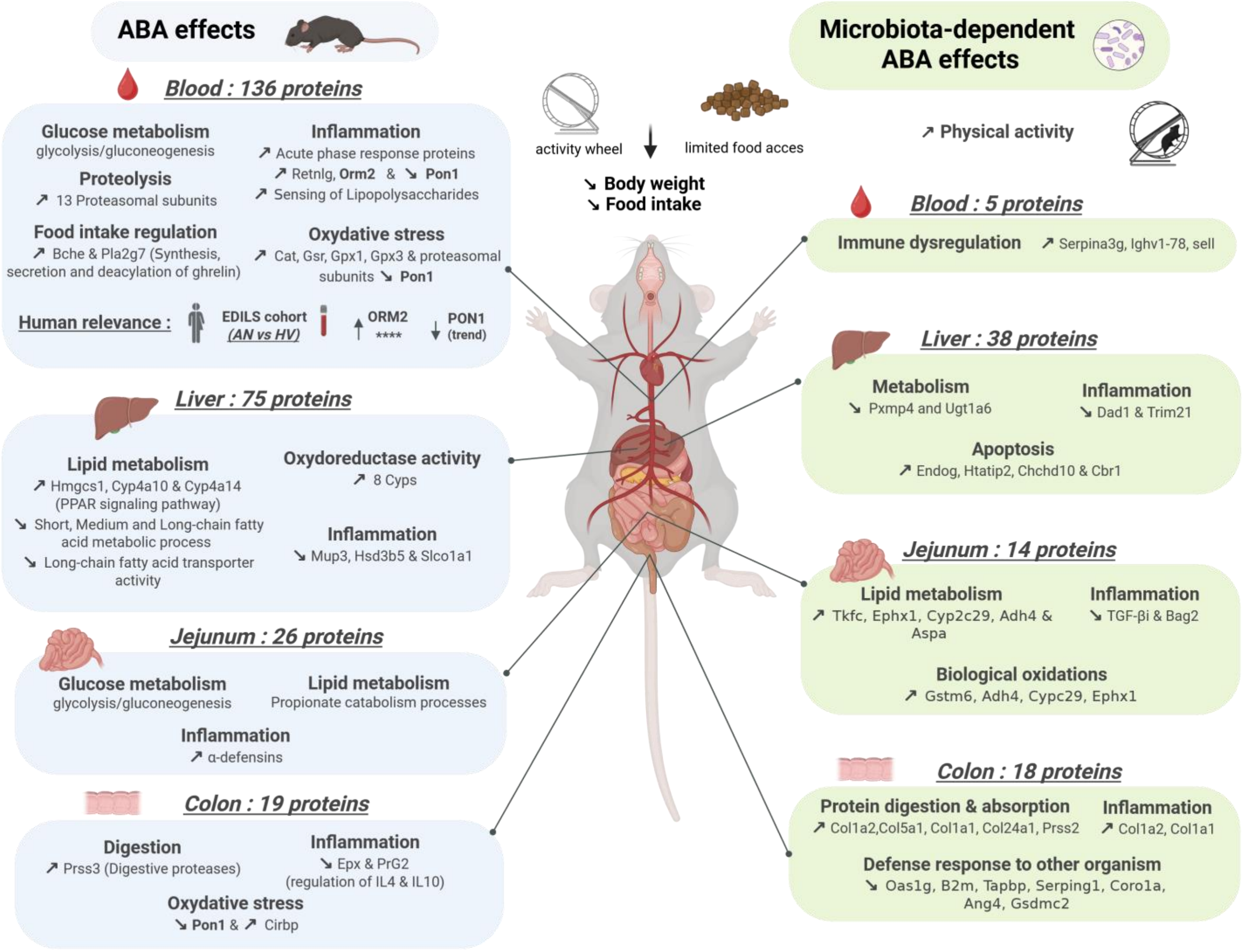

ABA effects
Blood : 136 proteins
Glucose metabolism
glycolysis/gluconeogenesis
Proteolysis
↗ 13 Proteasomal subunits
Food intake regulation
↗ Bche & Pla2g7 (Synthesis, secretion and deacylation of ghrelin)
Inflammation
↗ Acute phase response proteins
↗ Retnlg, Orm2 & ↘ Pon1
↗ Sensing of Lipopolysaccharides
Oxydative stress
↗ Cat, Gsr, Gpx1, Gpx3 & proteasomal subunits ↘ Pon1
Human relevance :
EDILS cohort (AN vs HV)
ORM2 ****
PON1 (trend)
Liver : 75 proteins
Lipid metabolism
↗ Hmgcs1, Cyp4a10 & Cyp4a14 (PPAR signaling pathway)
↘ Short, Medium and Long-chain fatty acid metabolic process
↘ Long-chain fatty acid transporter activity
Oxydoreductase activity
↗ 8 Cyps
Inflammation
↘ Mup3, Hsd3b5 & Slco1a1
Jejunum : 26 proteins
Glucose metabolism
glycolysis/gluconeogenesis
Lipid metabolism
Propionate catabolism processes
Inflammation
↗ α-defensins
Colon : 19 proteins
Digestion
↗ Prss3 (Digestive proteases)
Inflammation
↘ Epx & PrG2 (regulation of IL4 & IL10)
Oxydative stress
↘ Pon1 & ↗ Cirbp
activity wheel
limited food acces
↘ Body weight
↘ Food intake
Microbiota-dependent ABA effects
↗ Physical activity
Blood : 5 proteins
Immune dysregulation
↗ Serpina3g, Ighv1-78, sell
Liver : 38 proteins
Metabolism
↘ Pxmp4 and Ugt1a6
Inflammation
↘ Dad1 & Trim21
Apoptosis
↗ Endog, Htatip2, Chchd10 & Cbr1
Jejunum : 14 proteins
Lipid metabolism
↗ Tkfc, Ephx1, Cyp2c29, Adh4 & Aspa
Inflammation
↘ TGF-βi & Bag2
Biological oxidations
↗ Gstm6, Adh4, Cypc29, Ephx1
Colon : 18 proteins
Protein digestion & absorption
↗ Col1a2,Col5a1, Col1a1, Col24a1, Prss2
Inflammation
↗ Col1a2, Col1a1
Defense response to other organism
↘ Oas1g, B2m, Tapbp, Serping1, Corola, Ang4, Gsdmc2
